# The effect of COVID-19 mRNA vaccine on human lung carcinoma cells in vitro by means of Raman spectroscopy and imaging

**DOI:** 10.1101/2022.01.24.477476

**Authors:** Halina Abramczyk, Jakub Surmacki

## Abstract

The effect of COVID-19 mRNA vaccine on human lung carcinoma epithelial cells (A549) in vitro as a convenient preclinical model has been studied by means of Raman spectroscopy and imaging. The paper focuses on Raman imaging as a tool to explore apoptosis and oxidative phosphorylation in mitochondrial dysfunctions. The Raman results demonstrate alterations in the oxidation-reduction pathways associated with cytochrome c. We found that the COVID-19 mRNA vaccine downregulates the concentration of cytochrome c upon incubation with tumorous lung cells. Concentration of oxidized form of cytochrome c in mitochondria of lung cells decreases upon incubation the COVID-19 mRNA vaccine. Lower concentration of oxidized cytochrome c in mitochondria illustrates lower effectiveness of oxidative phosphorylation (respiration), reduced apoptosis and lessened ATP production. Moreover, mRNA vaccine increases significantly de novo lipids synthesis in lipid droplets up to 96 hours and alterations in biochemical composition. It seems that lipid composition of cells returns to the normal level for longer incubation time (14 days). In cell nucleus the mRNA vaccine does not produce statistically significant changes. The observed alterations in biochemical profiles upon incubation with mRNA vaccine in the specific organelles of the tumorous lung cells are similar to those we observe for other types of cancers, particularly brain glial cells.

## 1. Introduction

The pandemic outbreak in 2019 generating acute respiratory syndrome caused 494 587 638 confirmed cases of COVID-19, including 6 170 283 deaths, reported to WHO [1]. Pharmaceutical companies including Pfizer/BioNTech in 2020 prepared vaccines based on mRNA technology. The Pfizer/BioNTech vaccine (BNT162b2) was reported to be more than 90% effective against COVID-19 [2]. As of 5 April 2022, a total of 11 250 782 214 doses have been administered [1].

The pandemics has witnessed an explosion in research examining the interplay between the immune response and the intracellular metabolic pathways that mediate it. Research in the field of immunometabolism has revealed that similar mechanisms regulate the host response to infection, autoimmunity, and cancer. The new method of Raman imaging presented in this paper contributes to better understand pathways of our immune responses, recognize metabolites that regulates these pathways and suggest how to optimize mRNA technology to stimulate the adaptive immune system.

We will show that the key molecule in immunometabolism is cytochrome c. Cytochrome c plays a role as a key protein that is required in maintaining life (respiration) and cell death (apoptosis). Until now we do not know exactly if cytochrome c itself controls life (respiration) and death (apoptosis) mechanisms or rather these are other mechanisms caused by release of cytochrome c from the mitochondria to cytoplasm.

Dysregulation of oxidative phosphorylation and apoptosis in cells of the immune system can have essential consequences, which may result in diseases, including cancer and autoimmunity. This paper summarizes our current understanding of the role of cytochrome c in cancer and immune responses and aims to point out future directions of research.

In the paper, we study the effect of COVID-19 mRNA on the human carcinoma lung epithelial cells by using a novel non-invasive tool of Raman imaging. Here we demonstrate that Raman imaging gives new insight into basic mechanisms of cancer pathology and effect of mRNA vaccines on specific organelles in a single cell in vitro.

Raman spectroscopy and imaging provides quantitative and non-invasive monitoring of intra-cellular changes without the need of exogeneous labelling. Conventional methods of molecular biology require destroying of cell membranes to extract intracellular components for studying the biochemical changes inside specific organelles. In Raman imaging, we do not need to break cells to learn about biochemical composition of intracellular organelles. Visualization of alterations in biochemical composition in separate organelles is extremely valuable to monitor molecular mechanisms in cancer development and mechanisms of infections. Until now, no technology has proven effective for detecting concentration of specific chemical compounds in separate living cell organelles. Therefore, existing analytical technologies cannot detect the full extent of biolocalization of different chemical compounds inside and outside specific organelles.

We will study human cancer lung cells upon incubation with mRNA vaccine. The cancer diseases are the most serious cause of death, exceeding heart disease, strokes, pneumonia and COVID-19. There is no vaccine against most cancers, rapid development of mRNA vaccines may help in development anticancer vaccines in the near future.

The announcement of effective and safe vaccines for COVID-19 has been greeted with enthusiasm, but many fundamental mechanisms by which mRNA vaccines induce strong responses are still incompletely understood, particularly for patients suffering from cancer and studies should be continued [2]. It is not clear which cell specific activation contributes the most to vaccine efficacy and what mechanisms may inhibit adaptive immunity or lead to poor tolerability of the vaccine [4]. Possible short and long-term effects on cell metabolism from spike S protein produced by COVID-19 vaccination will be discussed. Researchers have warned that COVID-19 mRNA vaccine induces complex reprogramming of innate immune responses and biodistribution that should be considered in the development and use of mRNA-based vaccines [5,6]. It has been reported [6] that intramuscular vaccines in monkeys remain near the site of injection at the arm muscle and local lymph nodes. The lymphatic system also remove waste materials. Similar results were reported for a mRNA vaccine against H10N8 and H7N9 influenza viruses for animal model of mice [6]. However, recent results on interactions between the immune system and the viral proteins that induce immunity against COVID-19 suggest that the mechanisms may be more complex than previously expected [7]. Recently it has been reported that spike S protein of COVID-19 has remain not only near the site of injection, but also circulate in the blood. Clearance of detectable COVID-19 protein in blood plasma is correlated with production of IgG and IgA [8].

In the view of these results it is important to monitor biodistribution and location of metabolites upon mRNA vaccines in human host cells in vitro and appropriate experimental animal models. Visualization of chemical alterations in single cells upon delivery of mRNA vaccines would help evaluate the efficacy of candidate formulations and aid their rational design for preclinical and translational studies.

In this paper we will study the effect of COVID-19 mRNA vaccine on the human lung epithelial cancer cells in the respiratory system. Here, we will show that Raman imaging allows for quantitative, and non-invasive monitoring biochemical alterations in specific organelles in response to mRNA vaccine. We will compare the effect of mRNA on cancer lung cells with the effect of cancer itself (control). According to our best knowledge the mRNA vaccine has not been clinically tested for cancer patients. Therefore, this contribution will help monitoring responses in host lung cells similar to a viral infection, because the incubation with COVID-19 mRNA vaccine mimics some mechanisms of COVID-19 infection. Of course it has to be remembered that instead of the whole virus, only one protein S, essential for the immune response, is injected without COVID-19 virus replications.

We will study human lung cancer cells in vitro (A549) incubated with the COVID-19 mRNA vaccine by Raman imaging. We will monitor the effect of the mRNA vaccine on biodistribution of different chemical components, particularly cytochrome c, in the specific organelles of a cell: nucleus, mitochondria, lipid droplets, cytoplasm and membrane. In this paper we will explore alterations in reduction-oxidation pathways related to cytochrome c in human cancer lung cells upon incubation in vitro with COVID-19 mRNA vaccine.

We will demonstrate that Raman spectroscopy and Raman imaging are competitive clinical diagnostics tools for cancer diseases linked to mitochondrial dysfunction and are a prerequisite for successful pharmacotherapy of cancer. The strength of our approach is that data on the biology of pulmonary cells is of interest. Patients recovering from COVID-19 sometimes suffer from a vague syndrome of the respiratory system.

## 2. Materials and Methods

### 2.1. Reagents

Cytochrome c (C 2506) was purchased from Sigma Aldrich (Poland).

### 2.2 Cell culture, incubation with vaccine and preparation for Raman imaging

The human lung carcinoma cell line A549 (ATCC CCL-185) was purchased from American Type Culture Collection (ATCC). The A549 cells were maintained in F-12K medium (ATCC 30-2004) supplemented with 10% fetal bovine serum (ATCC 30-2020) without antibiotics in a humidified incubator at 37°C and 5% CO_2_ atmosphere. For Raman imaging cells were seeded on CaF_2_ window (Crystran Ltd, CaF_2_ Raman grade optically polished window 25mm dia x 1mm thick, no. CAFP25-1R) in 35 mm Petri dish at a density of 5×10^4^ cells per Petri dish. On the following day the culture medium was replaced with the culture medium (F-12K medium (ATCC 30-2004) supplemented with 10% fetal bovine serum) supplemented with 1 or 60 μL diluted COVID-19 mRNA vaccine per 1 mL of medium. We diluted the COVID-19 mRNA vaccine in 1.8 mL of 0.9% sodium chloride [9]. The real dose of the diluted vaccine that is administered to patients is equal to 0.3 mL corresponding to 30 μg mRNA per dose. In our experiments the total volume of used Petri dishes was 3 mL, so that give us doses of 0.3 and 18 μg mRNA per Petri dish (1 or 60 μL diluted vaccine per 1 mL of medium). The incubation time with the vaccine was 1, 24, 72, 96 hours and 14 days, respectively.

Before Raman examination, the growing medium was removed and the cells were fixed with 10% formalin for 10 minutes and kept in phosphate-buffered saline (PBS, Gibco no. 10010023) during the experiment.

### 2.3. Raman imaging and spectroscopy

Raman spectroscopy is an analytical method in which inelastic scattered light is used to obtain the information about the molecular vibrations. Raman scattering is an inelastic scattering process with a transfer of energy between the molecular vibrations (and rotations) and the scattered photons. If the molecule receive energy from the photon energy being excited to a higher vibrational, the scattered photon loses energy and the Stokes Raman scattering occurs. Inversely, if the molecule occupying higher vibrational state is excited and then loses energy by return to a lower vibrational level, the scattered photon gains the corresponding energy and Anti-Stokes Raman scattering occurs. The Stokes Raman scattering is always more intense than the Anti-Stokes component and for this reason we measured the Stokes signal by Raman spectroscopy. The Rayleigh, Stokes and Anti-Stokes Raman Scattering processes were presented in Scheme 1.

**Scheme 1.**
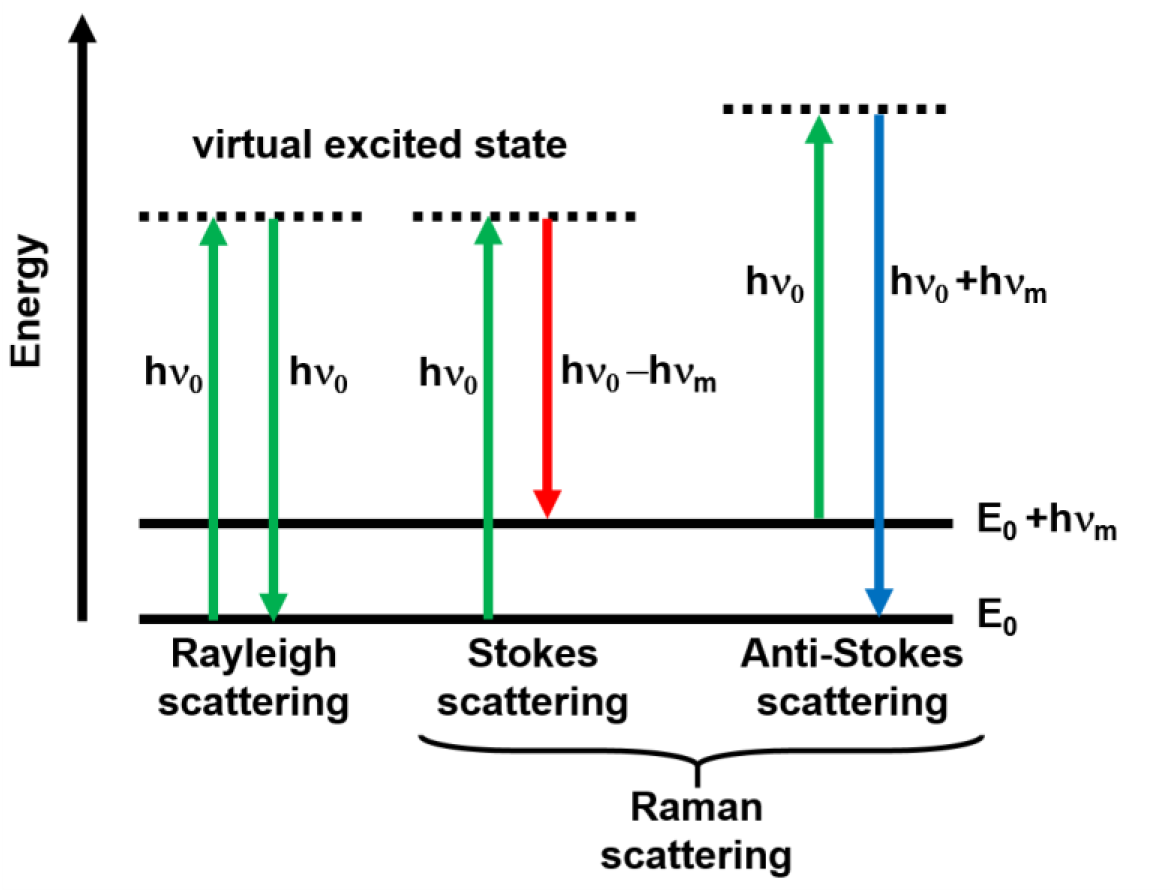
Schematic presentation of scattering phenomena.

Raman imaging is a technique based on Raman scattering allowing to measure vibrational spectra of any area. The imaging mode allows to analyze distribution of different chemical molecules inside the sample area. Using algorithms of an artificial intelligence such as Cluster Analysis (see section 2.4) based on 2D data make possible to create Raman maps to visualize cell’s organelles: nucleus, mitochondria, lipid structures, cytoplasm, cell membrane and learn about their biocomposition. Scheme 2 illustrates the idea of data acquisition for a single cell spectrum and Raman imaging.

**Scheme 2.**
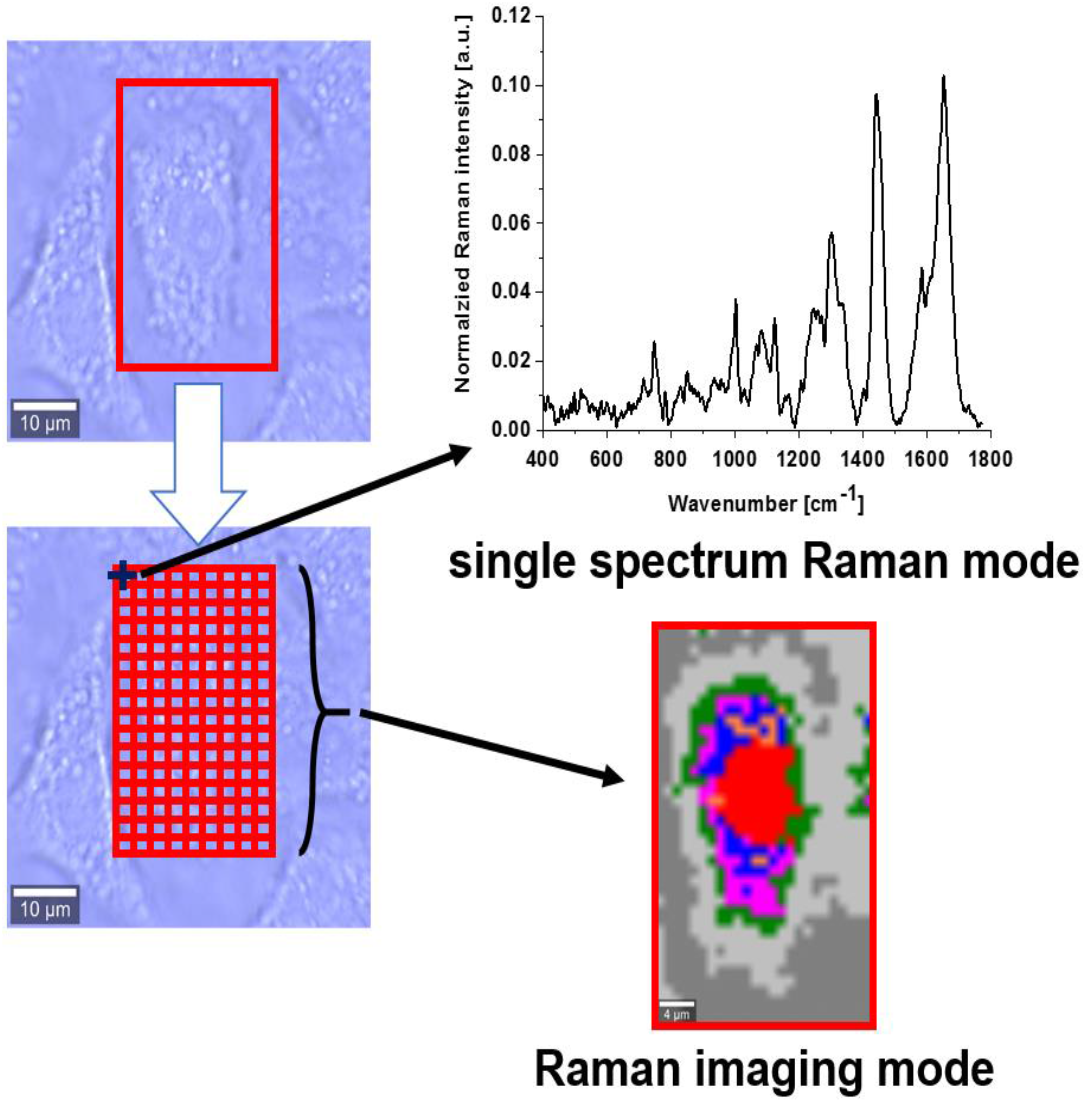
Data acquisition for a single point in a cell spectrum and Raman imaging of a whole cell (A549)

Raman spectra and images were recorded using a confocal Raman microscope (WITec (alpha 300 RSA+), Ulm, Germany) in the Laboratory of Laser Molecular Spectroscopy, Lodz University of Technology, Poland. The excitation laser at 532 nm was focused on the sample to the laser spot of 1 µm with the microscope (Olympus Düsseldorf, Germany) with A 40× water immersion objective (Zeiss, W Plan-Apochromat 40×/1.0 DIC M27 (FWD = 2.5 mm), VIS-IR) via an optical fiber with a diameter of 50 µm. The laser excitation power was 10 mW. Raman images were recorded with a spatial resolution of 1 × 1 µm and the collection time of 0.5 and 1 s for Raman images with a UHTS (Ultra-High-Throughput Screening) monochromator (WITec, Ulm, Germany) and a thermoelectrically cooled CCD camera ANDOR Newton DU970N-UVB-353 (EMCCD (Electron Multiplying Charge Coupled Device, Andor Technology, Belfast, Northern Ireland) chip with 1600 × 200 pixel format, 16 µm dimension each) at −60°C with full vertical binning. The Raman spectrometer was calibrated prior to the measurements using a silica plate with a maximum peak at 520.7 cm^-1^.

### 2.4. Data processing and cluster analysis

WITec Project Plus software was used to collect and process Raman data. To refine background subtraction and the normalization (model: divided by vector norm) we used Origin software. Spectroscopic data were analyzed using Cluster Analysis method. Details are given in our previous paper [10]. The normalization was performed for two regions: fingerprint region (370-1770 cm^-1^) and high frequency region (2700-3100 cm^-1^) separately. The Raman maps presented in the manuscript were constructed based on the number of clusters of 7. Each cluster is characterized by different average Raman spectrum and describe the inhomogeneous distribution of chemical components of different organelles within the cell.

## 3. Results

To properly address alterations in single lung cells upon incubation with COVID-19 mRNA vaccine, we systematically investigated how Raman spectroscopy and Raman imaging monitor responses to the vaccine in the specific organelles.

Figure 1 shows the Raman image of a typical cell of highly aggressive lung cancer incubated with mRNA vaccine for dose of 60 μL for the period of incubation of 96 hours and the Raman images of specific organelles. The Raman images were created by K-means cluster analysis (7 clusters). The blue, orange, magenta, red, green, light gray color represents lipids in lipid droplets and rough endoplasmic reticulum, lipid droplets filled with triacylglycerols of monounsaturated type (TAG), mitochondria, nucleus, cytoplasm, membrane, respectively. The dark grey color corresponds to area out of the cell.

**Figure 1.**
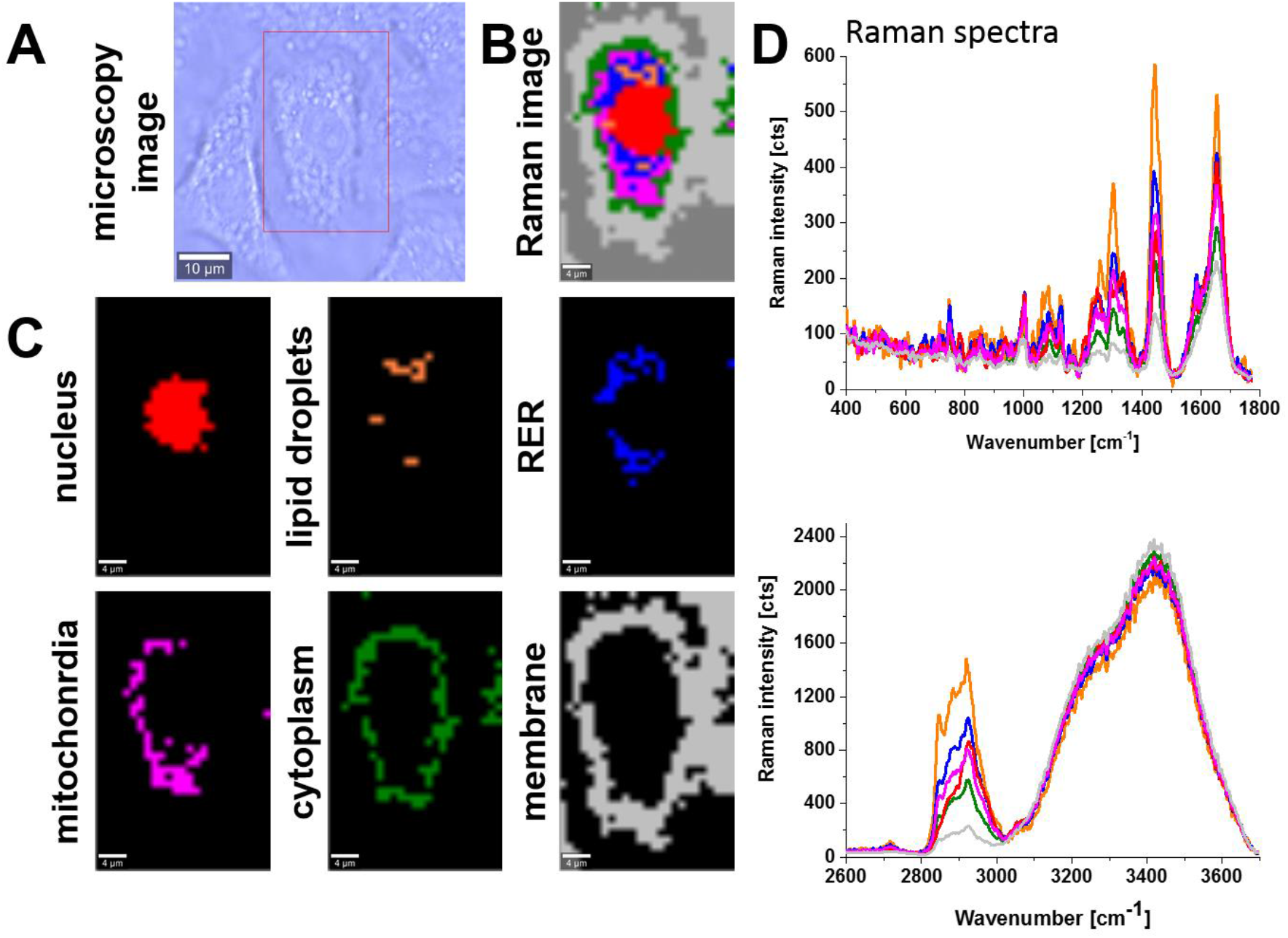
Microscopy image (A), Raman image of human lung carcinoma (A549) cell (25×40 μm, resolution 1.0 μm) incubated with COVID-19 mRNA vaccine (dose 60 μL) for 96 hours (B) and Raman images of specific organelles: nucleus (red), lipid droplets (orange), rough endoplasmic reticulum (RER, blue), mitochondria (magenta), cytoplasm (green) and membrane (light grey) (C) with corresponding Raman spectra (D) at 532 nm.

To track alterations in human lung cancer cells upon incubation with COVID-19 vaccine we systematically investigated how the average Raman spectra in each organelle of cells incubated with mRNA and control cells without mRNA incubation.

### 3.1. Mitochondria-mRNA

Figure 2 shows the effect of the COVID-19 mRNA vaccine on mitochondria in human lung carcinoma (A549) cell upon incubation for 1, 24, 72, 96 hours and 14 days for 1 μL and 60 μL at 1584, 1444 and 1654 cm^-1^.

**Figure 2.**
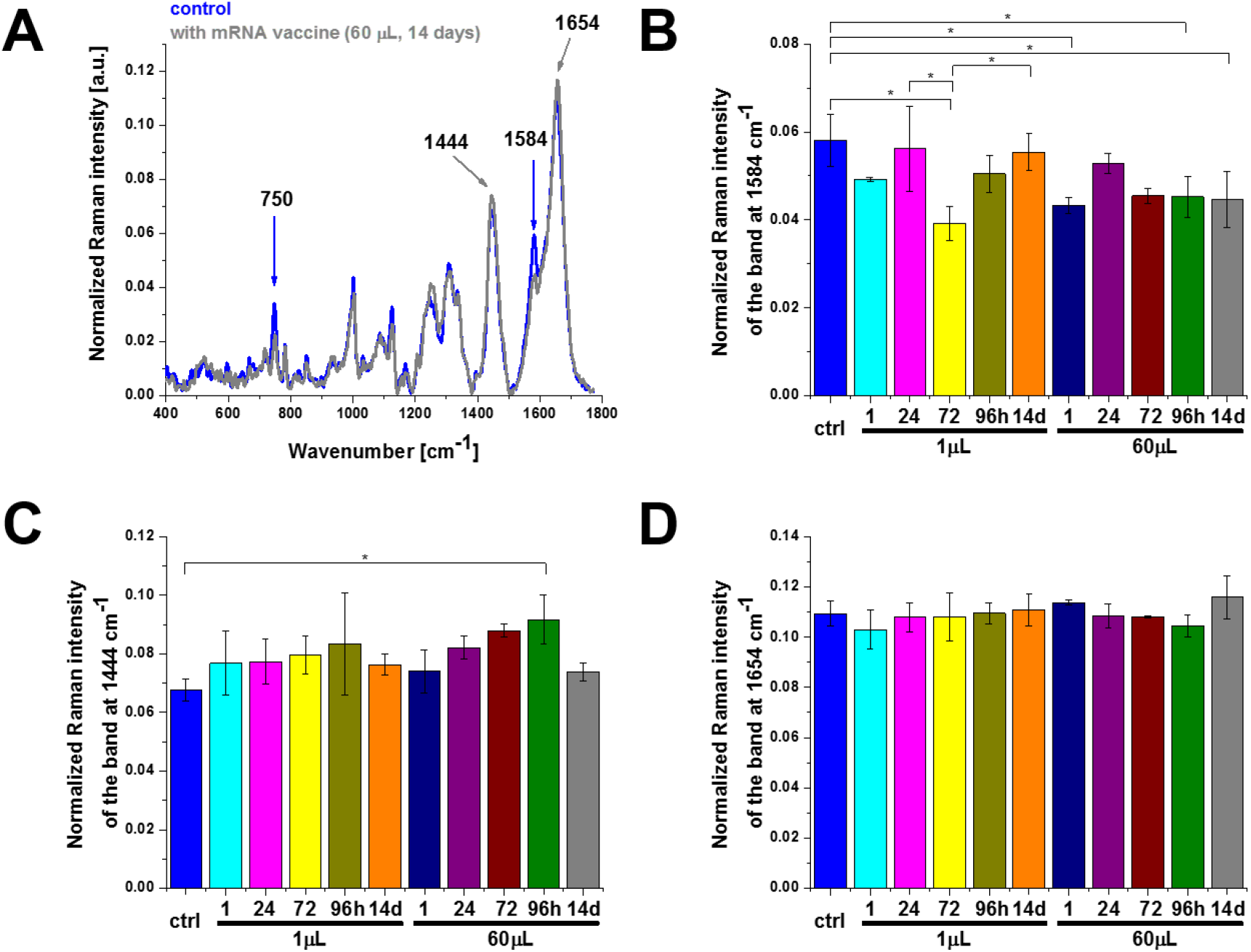
The effect of the COVID-19 mRNA vaccine on mitochondria in human lung carcinoma (A549) cell (A) upon incubation for 1, 24, 72, 96 hours and 14 days for 1 μL and 60 μL at 1584 (B), 1444 (C) and 1654 cm^-1^ (D) (number of cells: 3, number of Raman spectra for each single cell: minimum 1600, excitation wavelength: 532 nm, laser power: 10 mW, integration time: 1.0 sec). The statistical significance was calculated with the one-way ANOVA using the Tukey test, asterisk * denotes that the differences are statistically significant, p-value ≤ 0.05.

Detailed inspection into Figure 2 demonstrates that the most significant changes occur at 750 cm^-1^ and 1584 cm^-1^ corresponding to cytochrome c [11] and at 1444 cm^-1^ corresponding to saturated C-H bending vibrations of lipids [12]. Let us first concentrate on cytochrome c.

Briefly, there is a broad family of cytochromes that are classified on the basis of their lowest electronic energy absorption band in their reduced state, such as cytochrome P450 (450 nm), cytochrome c (Q band 550 nm), cytochromes b (≈565 nm), cytochromes a (605 nm). We used the laser excitation with the wavelength at 532 nm, whose energy is in coincidence with the electron absorption of the Q band. This resonance leads to greatly enhanced intensity of the Raman scattering which is called Resonance Raman Effect (RRE). RRE amplification facilitates the study of cytochrome c which is present at low concentrations under physiological conditions of humans. The cytochrome c is localized at the internal mitochondrial membranes between the complex known as complex III (sometimes called also Coenzyme Q – Cyt C reductase or cytochrome bc1 complex) and the complex IV. Cytochrome c plays a key role in the electron transport chain of the oxidative phosphorylation (respiration) process. The oxidized cytochrome c (cyt c Fe^3+^) is reduced to cytochrome c (cyt c Fe^2+^) by the electron obtained from the complex III. The reduced cytochrome c passes an electron to the copper binuclear center of the complex IV, being oxidized back to cytochrome c (cyt c Fe^3+^). The Complex IV, which is the final complex in the electron transport chain contains two cytochromes a and a3 and two copper centers. The electron transport chain is followed by a ATP synthase, which is often called complex V in the electron transport chain. Therefore, under normal conditions cytochrome c in mitochondrion exists in the oxidized form. Under pathological conditions metabolic processes in mitochondrion results in down-regulation of the transfer between cytochrome c and the complex IV leading to increase of the reduced form of cytochrome. Raman signals of the oxidized form are significantly lower than those of the reduced form of cytochrome c [10]. Therefore, Raman spectra can be used as markers of redox status of cytochrome c.

The peak at 1584 cm^-1^ in Figure 2 represents the “redox state Raman marker” of cytochrome c. Recently we demonstrated that this Raman vibration can serve as a sensitive indicator of oxidized and reduced forms of cytochrome c [10,11,13]. It indicates that the Raman peak at 1584 cm^-1^ can be used as a marker to explore apoptosis and oxidative phosphorylation in mitochondria [16,18]. This band reflects the dual activity of cytochrome c in life and death processes: apoptosis and oxidative phosphorylation. The balance between proliferation of cancer cells (oxidative phosphorylation) and cells death (apoptosis) decide about rate of cancer development and cancer aggressiveness [14,15]. The Raman signal of a single cell at 1584 cm^-1^ depends on cytochrome c concentration (which also depends on number of mitochondria in a cell), and the redox state (oxidized or reduced forms). The Raman intensity of the oxidized form is much weaker than that of the reduced form [10,13,14,16].

Inside a normal mitochondrion cytochrome c exists in the oxidized form. Dysfunction of mitochondrion associated with several malignancies, including cancer or virus infections blocks the transfer of electrons between complexes III and IV of the electron transport chain, resulting in lower efficiency of the oxidative phosphorylation (respiration) process and lower ATP synthesis. This results in change of the redox status of cytochrome c from the oxidized (cyt c Fe^3+^) to reduced form (cyt c Fe^2+^) [10]. Cytochrome c acts not only as mitochondrial charge carrier that transfers electrons between the complexes III and IV, but also activates the caspase cascade, when cytochrome c is released from the intermembrane space of mitochondrion to cytoplasm. The released cytochrome c interacts with apoptosis-protease activating factor 1 (Apaf-1) triggering the caspase cascade in the cell and induces an inflammatory response in the immune system. The activated caspases ultimately lead to cell apoptosis. Therefore, it was suggested that cytochrome c may play the role of an universal DAMP molecule (damage-associated molecular patterns) that informs the immune system of infection danger in cells or tissues [17]. DAMP molecules that is released from damaged or death cells interact with pattern recognition receptors (PRRs) and activate the innate immune system by [18]. Therefore cytochrome c plays a double function: it warns the immune system and contribute to the host’s defence, but also promote pathological inflammatory responses. Controlling cell death by apoptosis is necessary for the normal functioning of the immune system as well as is very important in natural mechanisms protecting against cancer. When less cytochrome c is released as DAMP molecules from the damaged or dying cells the innate immune system is not activated to the sufficient level to protect the body by interacting with PRRs.

The similar mechanism may play a key role in adaptive immune system, because it is reported that cytokines play an important role in adaptive immunity [20,21]. CD4^+^ helper T cells (Th) can be divided into two subgroups, type I helper T lymphocytes (Thl) and type II helper T cells (Th2) [20].

We provided evidence that the balance between apoptosis and the oxidative phosphorylation is regulated by interactions between cytochrome c and cardiolipin [10]. Cardiolipin-bound Cyt c, probably does not participate in electron shuttling of the respiratory chain [22] leading to production of reduced form of cytochrome c resulting in lower efficiency of respiration (oxidative phosphorylation) and lessened ATP production. The reduced form of cytochrome bound to cardiolipin cannot induce caspase and apoptosis process [10,22].

The intensity of the Raman signal at 1584 cm^-1^ corresponding to the concentration of cytochrome c in mitochondria depends on three factors: a) redox status (oxidized or reduced form of cytochrome c, b) release of cytochrome c to cytoplasm., c) number of cytochrome c molecules in mitochondria that depends on the number of mitochondria in a single cell, As the Raman signals of the reduced form are significantly higher than those of the oxidized form of cytochrome c [10] one can state that the Raman intensity of the band at 1584 cm^-1^ in Figure 2 A and B represents an oxidized form of cytochrome (cyt c Fe^3+^) both for the control and the cells upon incubation with the COVID-19 mRNA vaccine. In the next paragraph we will demonstrate that there is no release of cytochrome c to cytoplasm. It indicates that the concentration of cytochrome c (cyt c Fe^3+^) must be determined by the factor c): the number of mitochondria in a single cell. The results in Figure 2B show the Raman signal at 1584 cm^-1^ with the mRNA vaccine decreases when compared with the control (without the mRNA vaccine -blue color) indicating that the number of mitochondria in a single cell decrease upon incubation with mRNA, which results in reduced metabolic activities. The number of mitochondria in a cell depends upon the metabolic demands and may change from a few mitochondria to thousands of this organelle [23].

In the view of the results presented in Figure 2 one can state that mRNA vaccine does not block the transfer of electrons between complexes III and IV of the respiratory chain, but decrease the number of mitochondria in cells resulting in lower efficiency of oxidative phosphorylation and ATP synthesis.

It is worth of noticing that a similar effect of down regulation of cytochrome c concentration was reported for brain cancer cells vs. cancer aggressiveness [9,15]. We showed that the Raman signal of the band at 1584 cm^-1^ related to the concentration of cytochrome c in mitochondria of a single cell in vitro decreases with brain tumor aggressiveness [11,16].

### 3.2. Cytoplasm-mRNA

In order to check if the observed lower concentration of cytochrome c in mitochondria is related to release of cytochrome c (apoptosis) to cytoplasm we studied the localization of cytochrome c in cytoplasm. Figure 3 shows the effect of the COVID-19 mRNA vaccine on the Raman spectra in cytoplasm of human lung carcinoma (A549) by Raman imaging compared with the control cells. Figure 3 shows a comparison of the average Raman for cytoplasm at 532 nm excitation with and without COVID-19 mRNA vaccine. One can see that that the Raman signal at 1584 cm^-1^ in cytoplasm (Figure 3B) is the same for control cells and those incubated with mRNA within statistical significance at p-value ≤ 0.05. It means that the lower concentration of cytochrome c in mitochondria is not related to the release to cytoplasm. In the view of this result from Figure 3B one can state that the mRNA does not activate additional apoptosis via cytochrome c release and does not act as a cytoplasmic apoptosis-triggering agent.

**Figure 3.**
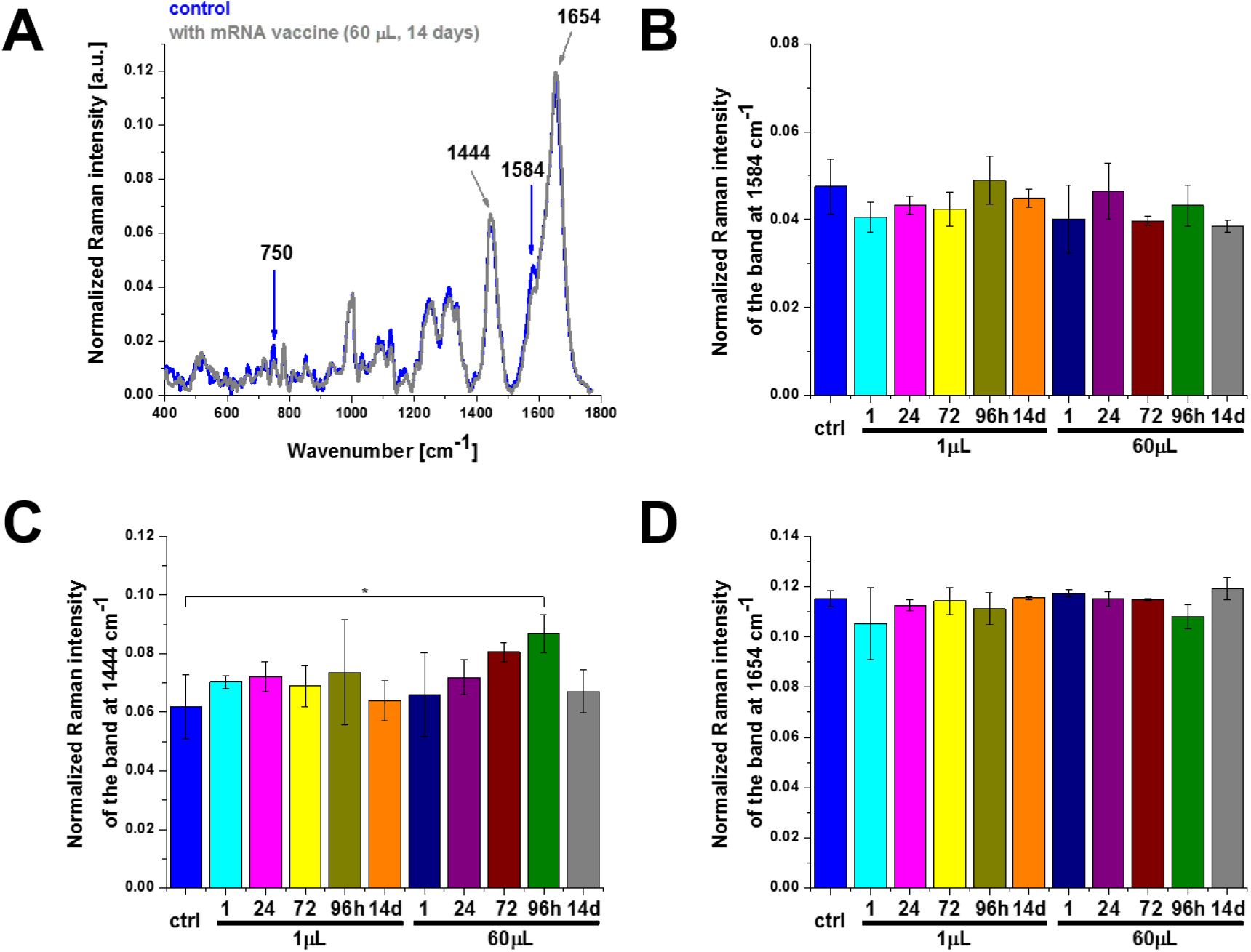
The effect of the COVID-19 mRNA vaccine on cytoplasm in human lung carcinoma (A549) cell upon incubation for 1, 24, 72, 96 hours and 14 days for 1 μL and 60 μL doses (number of cells: 3, number of Raman spectra for each single cell: minimum 1600, excitation wavelength: 532 nm, laser power: 10 mW, integration time: 1.0 sec). The one-way ANOVA using the Tukey test was used to calculate the value significance, asterisk * denotes that the differences are statistically significant, p-value ≤ 0.05.

### 3.3. Lipid droples-mRNA and rough endoplasmic reticulum-mRNA

The significant changes in cytochrome c concentration are also observed in other organelles. Similar as in mitochondria alterations in biochemical composition of cytochrome c reflected by the band at 1584 cm^-1^ are also observed in lipid droplets and in lipid structures of rough endoplasmic reticulum (ER) presented in Figures 4 and 5. Rough ER contains ribosomes that are responsible for protein synthesis (mRNA translation). Therefore, ribosomes are the sites where the protein synthesis for mRNA vaccines occurs. Ribosomes are too small to be seen by resolution of Raman imaging.

**Figure 4.**
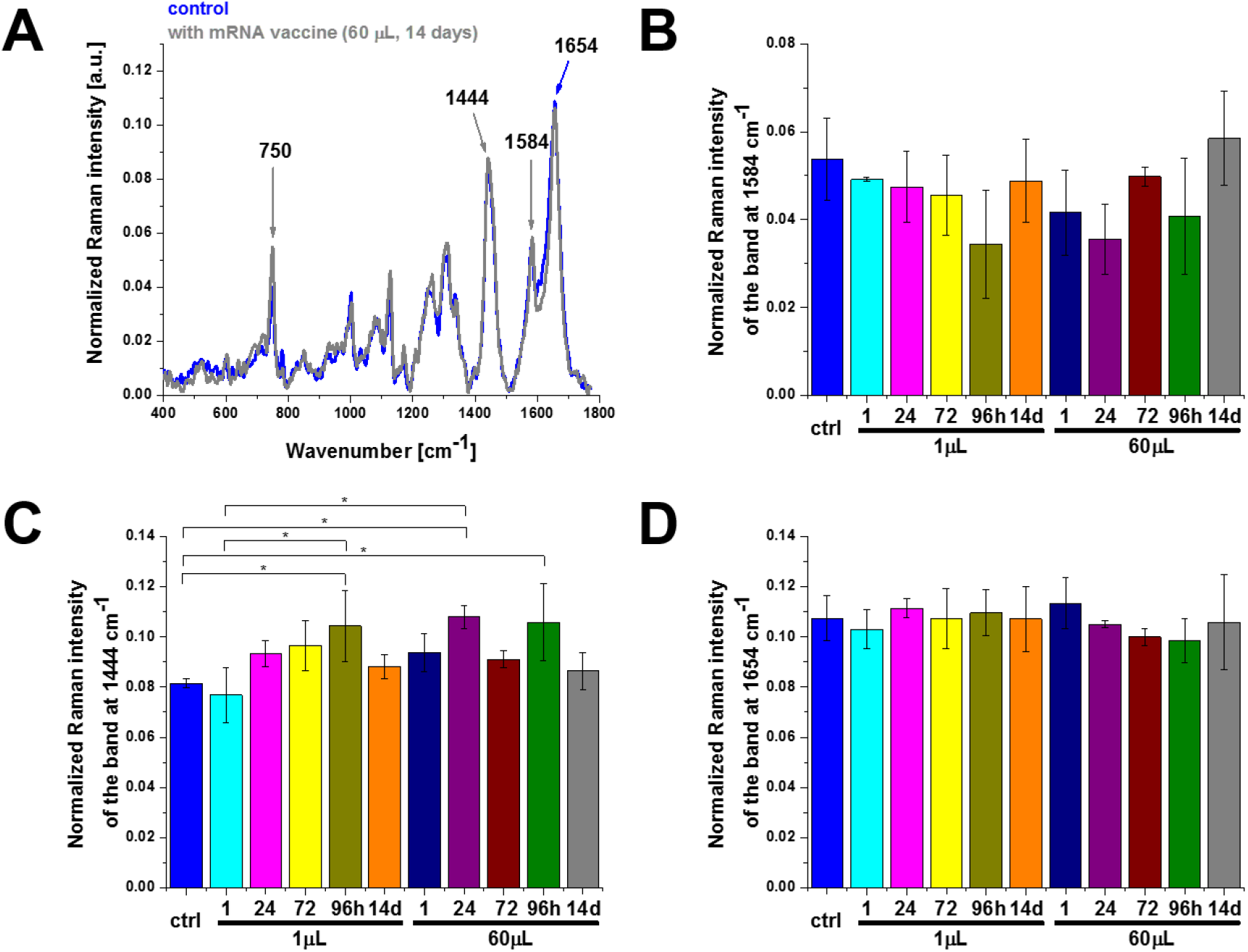
The effect of the COVID-19 mRNA vaccine on lipid droplets in human lung carcinoma (A549) cell upon incubation for 1, 24, 72, 96 hours and 14 days for 1 μL and 60 μL doses (number of cells: 3, number of Raman spectra for each single cell: minimum 1600, excitation wavelength: 532 nm, laser power: 10 mW, integration time: 1.0 sec). The one-way ANOVA using the Tukey test was used to calculate the value significance, asterisk * denotes that the differences are statistically significant, p-value ≤ 0.05.

**Figure 5.**
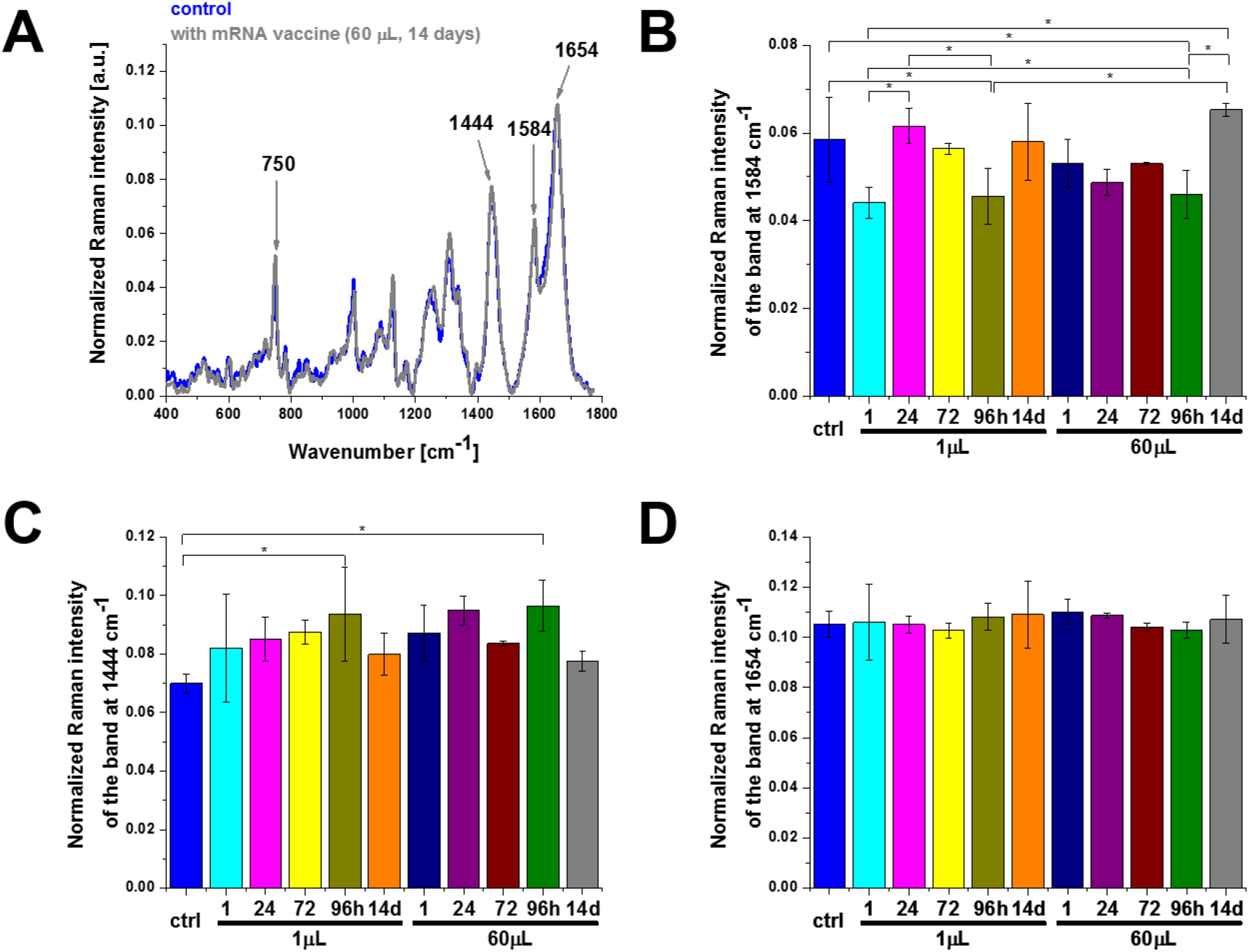
The effect of the COVID-19 mRNA vaccine on rough endoplasmic reticulum in human lung carcinoma (A549) cell upon incubation for 1, 24, 72, 96 hours and 14 days for 1 μL and 60 μL doses (number of cells: 3, number of Raman spectra for each single cell: minimum 1600, excitation wavelength: 532 nm, laser power: 10 mW, integration time: 1.0 sec). The one-way ANOVA using the Tukey test was used to calculate the value significance, asterisk * denotes that the differences are statistically significant, p-value ≤ 0.05.

Now we will focus on lipid synthesis upon mRNA vaccine. Figures 2-5 show that the Raman bands at 1444 cm^-1^ are significantly modified upon incubation with mRNA. Indeed, the Raman signal at 1444 cm^-1^ corresponding to C-H bending vibrations of lipids in lipid droplets (orange color in Figure 1) and in rough endoplasmic reticulum (blue color in Figure 1) clearly increases upon incubation with COVID-19 mRNA vaccine. The results indicate that mRNA vaccine up-regulates de novo lipid synthesis up to 96 hours. Similar trend was found for lipid reprogramming in cancers as a function of aggressiveness [24].

### 3.4. Nucleus-mRNA

It is claimed that mRNA vaccine does not enter the nucleus of the cell where our DNA (genetic material) is kept [4]. The explanation is the following: 1) The cell breaks down and gets rid of the mRNA soon after it finished translation procedure, 2) neither the coronavirus nor the RNA vaccines have a reverse transcriptase. Therefore, RNA vaccines cannot produce DNA molecules, 3) the cells keep their compartments well separated, and messenger RNAs cannot travel from the cytoplasm to the nucleus, because the life time of mRNA molecules is short. The mRNA as short-lived molecules and are degraded after a few hours, 4) reported clinical studies have shown no sign of DNA modification so far. These facts suggest that the vaccines do not alter our genome. Figure 6 shows results obtained for nucleus without and upon incubation with vaccine by Raman imaging. Our results support the conclusions that mRNA vaccine does not enter the DNA of the cell as we observe no statistically significant changes in the Raman signals.

**Figure 6.**
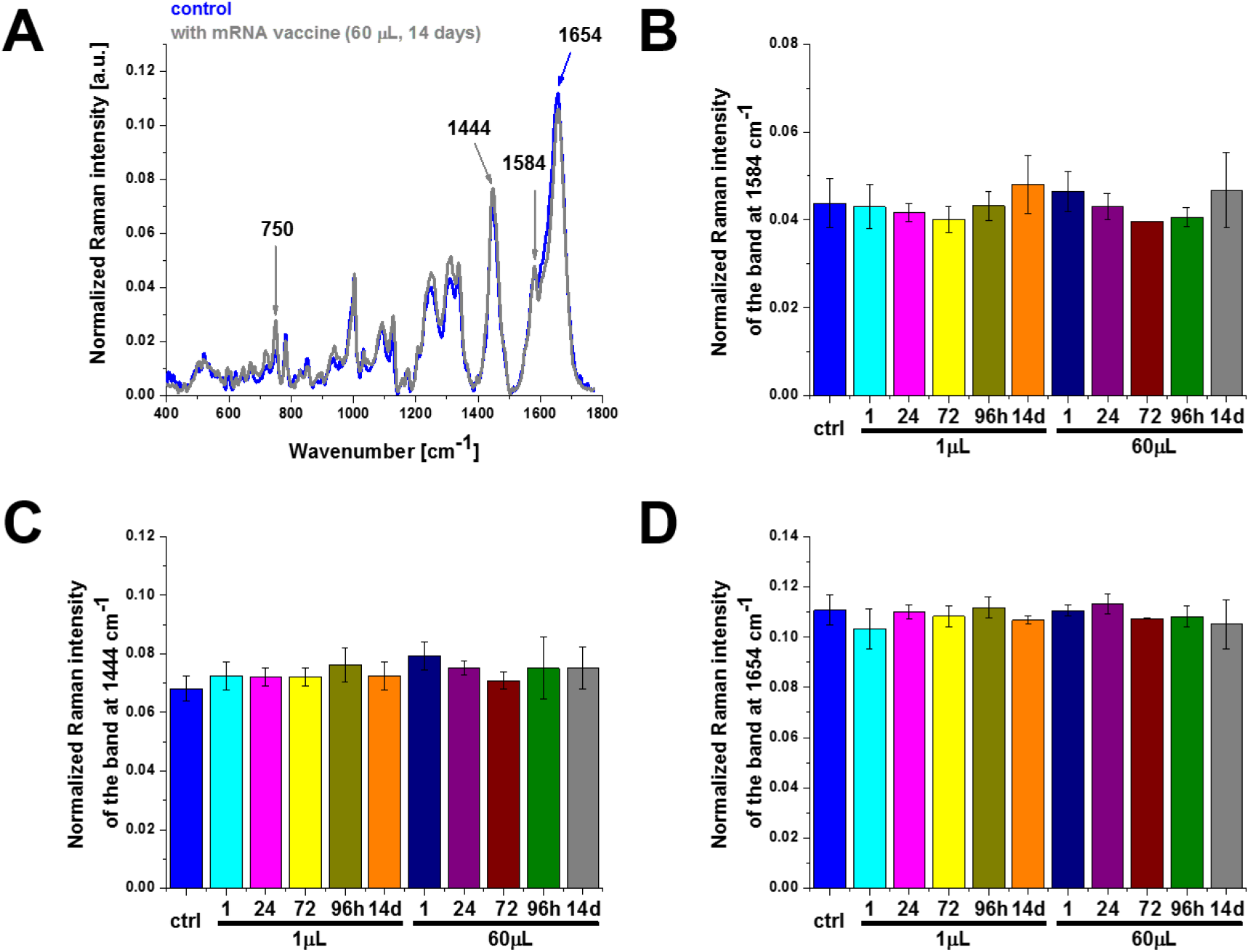
The effect of the COVID-19 mRNA vaccine on nucleus in human lung carcinoma (A549) cell upon incubation for 1, 24, 72, 96 hours and 14 days for 1 μL and 60 μL doses (number of cells: 3, number of Raman spectra for each single cell: minimum 1600, excitation wavelength: 532 nm, laser power: 10 mW, integration time: 1.0 sec). The one-way ANOVA using the Tukey test was used to calculate the value significance, asterisk * denotes that the differences are statistically significant, p-value ≤ 0.05.

### 3.5. Membrane-mRNA

As it is well known professional antigen-presenting cells (APCs) play a crucial role in initiating immune responses and are mainly represented by professional APC dendritic cells. However, under pathological conditions also epithelial cells act as nonprofessional APCs, thereby regulating immune responses at the site of exposure. Therefore, it is interesting to monitor alterations at the surface of the cell membranes of the lung epithelial cells upon incubation with mRNA.

Figure 7 shows effect of incubation with COVID-19 mRNA vaccine compared to the control cells without mRNA at the surface of the membrane. One can see that cytochrome c activity at 1584 cm^-1^ does not change with statistical significance at p-value ≤ 0.05 and also the Raman signals at 1444 cm^-1^ and 1654 cm^-1^ do not change within statistical significance.

**Figure 7.**
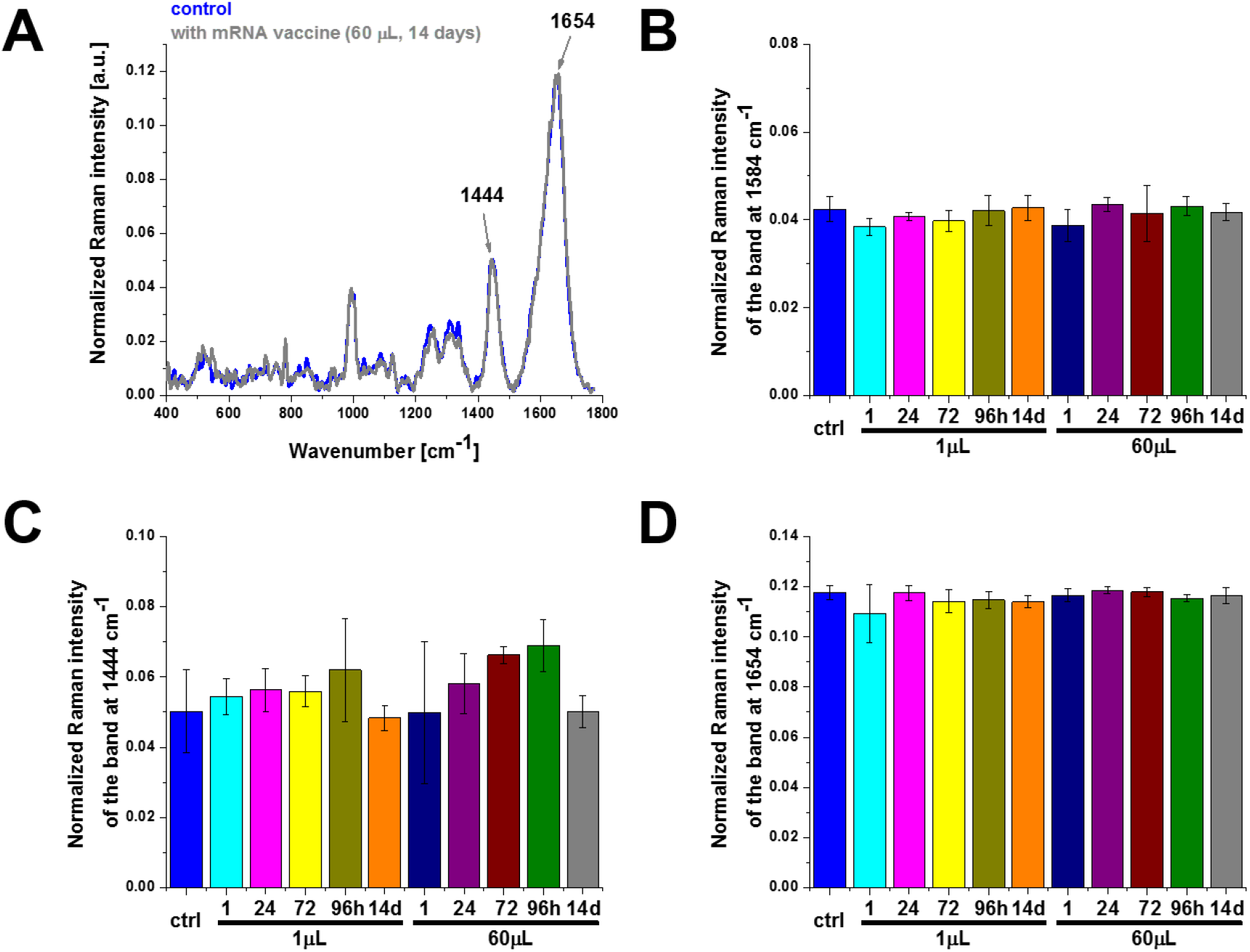
The effect of the COVID-19 mRNA vaccine on membrane in human lung carcinoma (A549) cell upon incubation for 1, 24, 72, 96 hours and 14 days for 1 μL and 60 μL doses (number of cells: 3, number of Raman spectra for each single cell: minimum 1600, excitation wavelength: 532 nm, laser power: 10 mW, integration time: 1.0 sec). The one-way ANOVA using the Tukey test was used to calculate the value significance, asterisk * denotes that the differences are statistically significant, p-value ≤ 0.05.

## 4. Conclusions

The effect of COVID-19 mRNA vaccine on human lung carcinoma epithelial cells (A549) in vitro has been studied by means of Raman spectroscopy and imaging. We studied biodistribution of different chemical components, particularly cytochrome c and lipids in the specific organelles of the cells: mitochondria, nucleus, lipid droplets, rough endoplasmic reticulum, cytoplasm and membrane and we observed biochemical alterations upon incubation with COVID-19 mRNA vaccine.

Cytochrome c activity in the efficiency of oxidative phosphorylation and apoptosis, is down-regulated by the COVID-19 mRNA vaccine. Lower concentration of oxidized cytochrome c observed in mitochondria in human lung cancer cells upon incubation with COVID-19 mRNA vaccine leads to reduced oxidative phosphorylation (respiration), and lessened ATP production. Incubation in vitro cells with mRNA vaccine increases significantly de novo lipids synthesis in lipid droplets and rough endoplasmic reticulum. For longer incubation time (14 days) it seems that lipid composition of cells returns to the normal level. mRNA vaccine does not produce statistically significant changes in the nucleus.

## Author Contributions

Conceptualization—H.A.; investigation —J.S.; visualization— J.S. and H.A.; writing—original draft — H.A.; writing—review and editing— J.S. and H.A. All authors have read and agreed to the published version of the manuscript.

## Institutional Review Board Statement

Not applicable.

## Informed Consent Statement

Not applicable.

## Data Availability Statement

The raw data underlying the results presented in the study are available from Lodz University of Technology Institutional Data Access. Request for access to those data should be addressed to the Head of Laboratory of Laser Molecular Spectroscopy, Institute of Applied Radiation Chemistry, Lodz University of Technology. Data requests might be sent by email to the secretary of the Institute of Applied Radiation Chemistry.

